# A global cancer data integrator reveals principles of synthetic lethality, sex disparity and immunotherapy

**DOI:** 10.1101/2021.01.08.425918

**Authors:** Christopher Yogodzinski, Abolfazl Arab, Justin R. Pritchard, Hani Goodarzi, Luke A. Gilbert

## Abstract

Advances in cancer biology are increasingly dependent on integration of heterogeneous datasets. Large scale efforts have systematically mapped many aspects of cancer cell biology; however, it remains challenging for individual scientists to effectively integrate and understand this data. We have developed a new data retrieval and indexing framework that allows us to integrate publicly available data from different sources and to combine publicly available data with new or bespoke datasets. Beyond a database search, our approach empowered testable hypotheses of new synthetic lethal gene pairs, genes associated with sex disparity, and immunotherapy targets in cancer. Our approach is straightforward to implement, well documented and is continuously updated which should enable individual users to take full advantage of efforts to map cancer cell biology.

## Introduction

Large scale but often independent efforts have mapped phenotypic characteristics of more than one thousand human cancer cell lines. Despite this, static lists of univariate data generally cannot identify the underlying molecular mechanisms driving a complex phenotype.

We hypothesized that a global cancer data integrator that could incorporate many types of publicly available data including functional genomics, whole genome sequencing, exome sequencing, RNA expression data, protein mass spectrometry, DNA methylation profiling, ChIP-seq, ATAC-seq, and metabolomics data would enable us to link disease features to gene products ^1–15^. We set out to build a resource that enables cross platform correlation analysis of multi-omic data as this analysis is in and of itself is a high-resolution phenotype. Multi-omic analysis of functional genomics data with genomic, metabolomic or transcriptomic profiling can link cell state or specific signaling pathways to gene function ^2,3,13,15–18^. Lastly, co-essentiality profiling across large panels of cell lines has revealed protein complexes and co-essential modules that can assign function to uncharacterized genes ^19^.

Problematically, in many cases publicly available data are poorly integrated when considering information on all genes across different types of data and the existing data portals are inflexible. For example, lists of genes cannot be queried against groups of cell lines stratified by mutation status or disease subtype. Furthermore, one cannot integrate new data derived from individual labs or other consortia.

We created the Cancer Data Integrator (CanDI) which is a series of python modules designed to seamlessly integrate genomic, functional genomic, RNA, protein and metabolomic data into one ecosystem. Our python framework operates like a relational database without the overhead of running MySQL or Postgres and enables individual users to easily query this vast dataset and add new data in flexible ways. This was achieved by unifying the indices of these datasets via index tables that are automatically accessed through CanDI’s biologically relevant Python Classes. We highlight the utility of CanDI through four types of analysis to demonstrate how complex queries can reveal previously unknown molecular mechanisms in synthetic lethality, sex disparity and immunotherapy. These data nominate new small molecule and immunotherapy anti-cancer strategies in KRAS-mutant colon, lung and pancreatic cancers.

## Results

### CanDI is a global cancer data integrator

We set out to integrate three types of data by creating programmatic and biologically relevant abstractions that allow for flexible cross referencing across all datasets. Data from the Cancer Cell Line Encyclopedia (CCLE) for RNA expression, DNA mutation, DNA copy number and chromosome fusions across more than 1000 cancer cells lines was integrated into our database with the functional genomics data from the Cancer Dependency Map (DepMap) (Fig. 1a,b and Supplementary Fig. 1) ^1,12,20^. We also integrated protein-protein interaction data from the CORUM database along with three additional distinct protein localization databases ^4,7,11,21^. CanDI by default will access the most recent release of data from DepMap although users can also specify both the release and data type that is accessed. The key advantage to this approach is that CanDI enables one to easily input user defined queries with multi-tiered conditional logic into this large integrated dataset to analyze gene function, gene expression, protein localization and protein-protein interactions.

**Figure 1.**
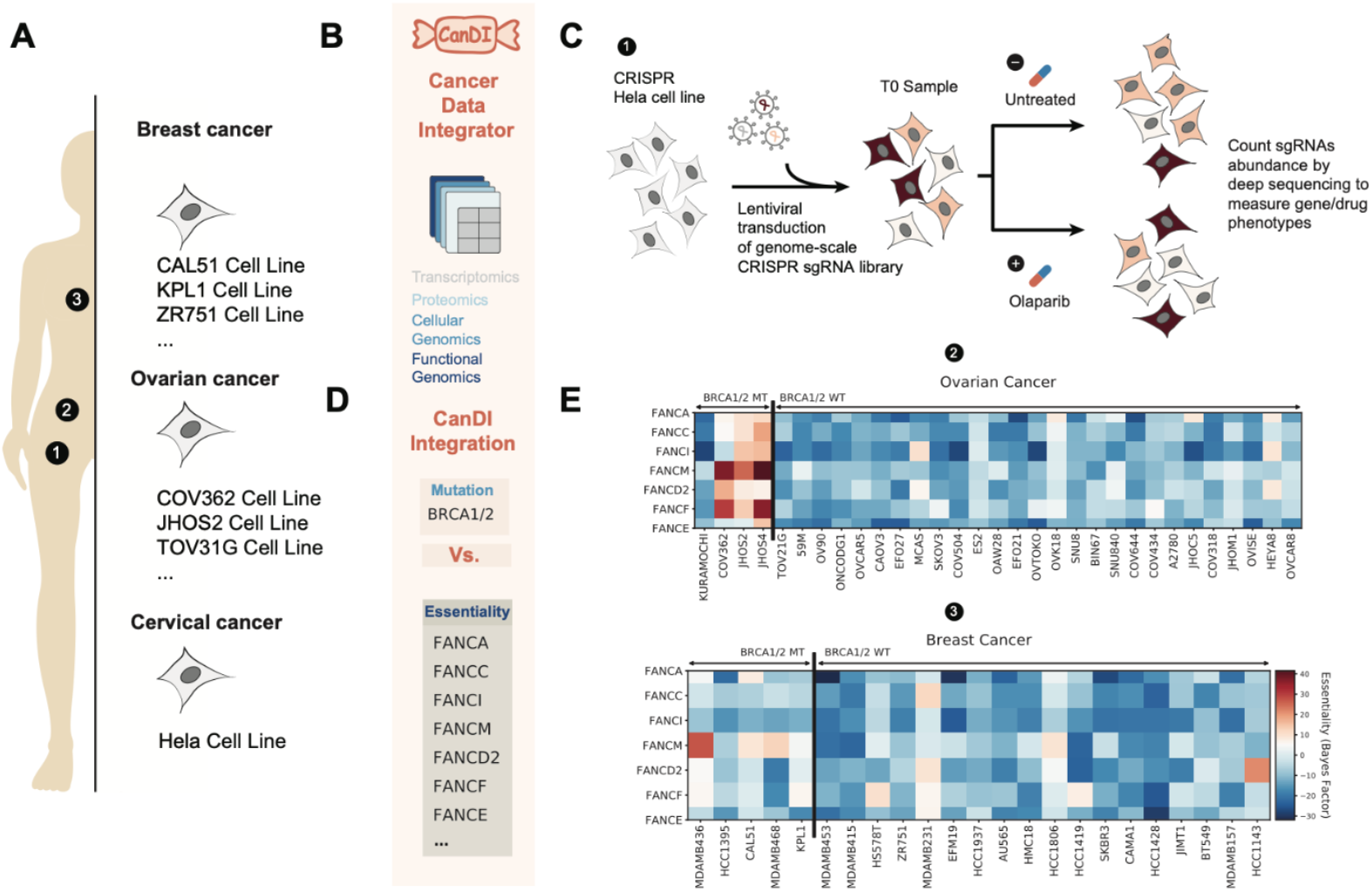
**(A)** A schematic showing human cell models integrated by CanDI. **(B)** A schematic illustrating types of data integrated by CanDI. **(C)** A cartoon of a genome-scale CRISPRi screen to identify genes that modulate response to PARP inhibition by Olaparib. **(D)** A schematic depicting data feature inputs parsed by CanDI. **(E)** Essentiality of Fanconi Anemia genes in ovarian and breast cancer cell lines separated by BRCA mutation status. A Bayes Factor score of gene essentiality is displayed by a heat map. N=4 BRCA1/2-mutant ovarian cancer, N=27 BRCA-wildtype ovarian cancer, N=5 BRCA1/2-mutant breast cancer, N=19 BRCA1/2-wildtype breast cancer.

### CanDI identifies genes that are conditionally essential in BRCA-mutant ovarian cancer

The concept that loss-of-function tumor suppressor gene mutations can render cancer cells critically reliant on the function of a second gene is known as synthetic lethality. Despite the promise of synthetic lethality, it has been challenging to predict or identify genes that are synthetic lethal with commonly mutated tumor suppressor genes. While there are many underlying reasons for this challenge, we reasoned that data integration through CanDI could identify synthetic lethal interactions missed by others.

A paradigmatic example of synthetic lethality emerged from the study of DNA damage repair (DDR)^22^. Somatic mutations in the DNA double-strand break (DSB) repair genes, *BRCA1/2*, create an increased dependence on DNA single strand break (SSB) repair. This dependence can be exploited through small molecule inhibition of PARP1 mediated SSB repair. Inhibition of PARP provides significant clinical responses in advanced breast and ovarian cancer patients but they ultimately progress^22^. Thus, new synthetic lethal associations with *BRCA1/2* are a potential path towards therapeutic development PARP refractory patients.

To illustrate the flexibility of CanDI to mine context specific synthetic sick lethal (SSL) genetic relationships we hypothesized that the genes that modulate response to a PARP1 inhibitor might be enriched for selectively essential proliferation or survival of BRCA1/2-mutant cancer cells. To test this hypothesis, we integrated the results of an existing CRISPR screen that identified genes that modulate response to the PARP inhibitor olaparib^23^. We then tested whether any of these genes are differentially essential for cell proliferation or survival in ovarian cancer and in breast cancer cell models that are either BRCA1/2 proficient or deficient (Fig. 1c,d). This query revealed that the Fanconi Anemia pathway is selectively essential in BRCA1/2-mutated ovarian cancer models but not in BRCA1/2-wild type ovarian cancer, BRCA1/2-mutated breast cancer or BRCA1/2-wildtype breast cancer models (Fig. 1e and Supplementary Table 1). To our knowledge a SSL phenotype between FANCM and BRCA1/2 has never been reported although a recent paper nominated a role for FANCM and BRCA1 in telomere maintenance^24^.

Importantly, FANCM is a helicase/translocase and thus considered to be a druggable target for cancer therapy^25^. Clinical genomics data support this SSL hypothesis although this remains to be tested in ovarian cancer patient samples^26^. Because the DepMap currently only allows single genes to be queried and does not enable users to easily stratify cell lines by mutation such analysis would normally take a user several days to complete manually. Our approach enabled this analysis to be completed using a desktop computer in less than two hours, which includes the visualization of data presented here (Fig. 1e).

### Conditional genetic essentiality in KRAS- and EGFR-mutant NSCLC cells

Beyond TSGs, many common driver oncogenes such as KRAS^G12D^ are currently undruggable, which motivates the search for oncogene specific conditional genetic dependencies.

We reasoned that CanDI enables us to rapidly search functional genomics data for genes that are conditionally essential in lung cancer cells driven by KRAS- and EGFR-mutations. We stratified non-small cell lung cancer cell (NSCLC) models by EGFR and KRAS mutations and then looked at the average gene essentiality for all genes within each of these 4 subtypes of NSCLC. We observed that KRAS is conditionally self-essential in KRAS-mutant cell models but that no other genes are conditionally essential in KRAS-mutant, EGFR-mutant, KRAS-wildtype or EGFR-wildtype cell models (Fig. 2a,b and Supplementary Table 2). This finding demonstrates that very few---if any---genes are synthetic lethal with KRAS- or EGFR-in KRAS- and EGFR-mutant lung cancer cell lines. It may be that these experiments are underpowered or it may be that when the genetic dependencies of diverse cell lines representing a disease subtype are averaged across a single variable (e.g. a KRAS-mutation) very few common synthetic lethal phenotypes are observed^27^. CanDI provides potential solutions for both of these hypotheses.

**Figure 2.**
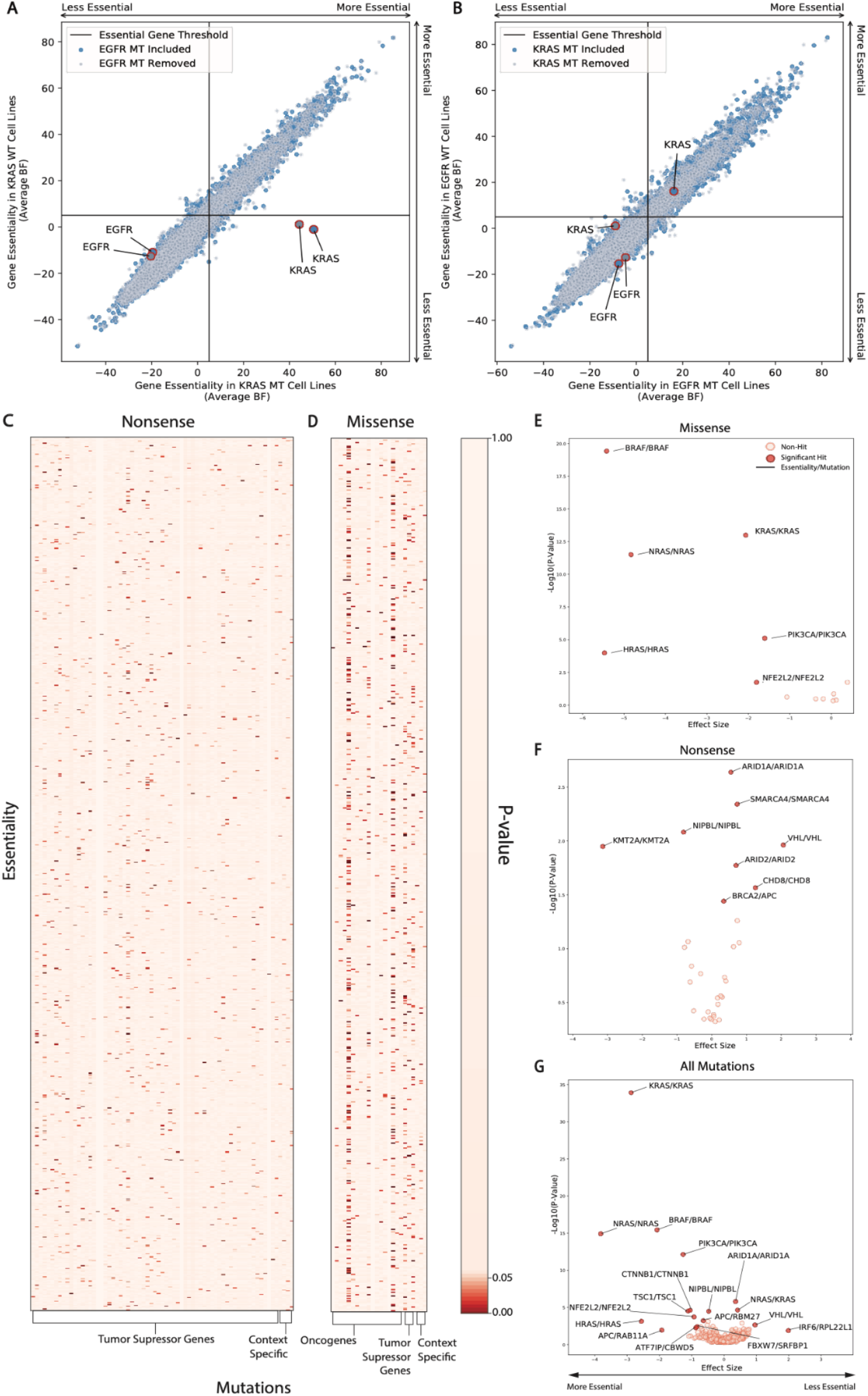
**(A)** Average gene essentiality for *KRAS* and *EGFR* in groups of NSCLC cell lines stratified by KRAS mutation status or by both KRAS and EGFR mutation status. N=38 for KRAS-wildtype shown in blue N=19 for KRAS-mutant shown in blue. N=30 for KRAS-wildtype EGFR-wildtype shown in grey and N=16 for KRAS-mutant EGFR-wildtype shown in grey. Gene essentiality is an averaged Bayes Factor score for each group of cell lines. **(B)** Average gene essentiality for *KRAS* and *EGFR* in groups of NSCLC cell lines stratified by EGFR mutation status or by both EGFR and KRAS mutation status. N=46 for EGFR-wildtype shown in blue, N=11 for EGFR-mutant shown in blue. N=30 for EGFR-wildtype KRAS-wildtype shown in grey and N=8 for EGFR-mutant KRAS-wildtype shown in grey. Gene essentiality is an averaged Bayes Factor score for each group of cell lines. **(C)** P-values from Chi^2^ tests of gene essentiality and nonsense mutations. **(D)** P-values from Chi^2^ tests of gene essentiality and missense mutations. **(E)** A scatter plot showing effect size of the change in gene essentiality with select missense mutations and the −Log10(P-value) of each essentiality/mutation pair. **(F)** A scatter plot showing effect size of the change in gene essentiality with select nonsense mutations and the −Log10(P-value) of each essentiality/mutation pair. **(G)** A scatter plot showing effect size of the change in gene essentiality with all mutations and the −Log10(P-value) of each essentiality/mutation pair.

### CanDI enables a global analysis of conditional essentiality in cancer

It is thought that data aggregation across vast landscapes of unknown co-variates does not necessarily increase the statistical power to identify rare associations^27^. Thus, the global analyses of aggregated cancer data sometimes lies in systematically sub setting data based on key co-variates post aggregation. This has been observed in driver gene identification^28^. Inspired by our analysis of TSG and oncogene conditionally essentiality above, we next used CanDI to identify genes that are conditionally essential in the context of several hundred cancer driver mutations.

We first grouped driver mutations (e.g. nonsense or missense) for each driver gene. For this analysis, we selected several thousand genes that are in the 85-90^th^ percentile of essentiality within the DepMap data and therefore conditionally essential, meaning these genes are required for cell growth or survival in a subset of cell lines. Importantly, it is not known why these several thousand genes are conditionally essential. We then tested whether each of these conditionally essential genes has a significant association with individual driver mutations. Our analytic approach does not weight the number of cell models representing each driver mutation nor does this give information on phenotype effect sizes. Our analysis nominates a large number of conditionally dependent genetic relationships with both TSG and oncogenes (Fig. 2c,d and Supplementary Table 3). A number of the conditional genetic dependencies identified in our independent variable analysis above are represented by a limited number of cell models and so further investigation is needed to validate these conditional dependencies, but this data further suggests that averaging genetic dependencies across diverse cell lines with un-modeled covariates obscures conditional SSL relationships.

To further investigate this hypothesis, we analyzed these same conditional genetic relationships with a second analytic approach that weights the number of cell models representing each driver mutation. We observed a limited number of conditional genetic dependencies that largely consists of oncogene self-essential dependencies as previously highlighted for KRAS-mutant cell lines (Fig. 2e-g and Supplementary Table 4)^13,29^. Thus, analysis that averages each conditional phenotype across diverse panels of cell lines with unknown covariates masks interesting conditional genetic dependencies.

### CanDI reveals female and male context specific essential genes in colon, lung and pancreatic cancer

Cancer functional genomics data is often analyzed without consideration for fundamental biological properties such as the sex of the tumor from which each cell line is derived. It is well established that biological sex influences cancer predisposition, cancer progression and response to therapy^30^. We hypothesized that individual genes may be differentially essential across male and female cell lines. This hypothesis to our knowledge has never been tested in an unbiased large-scale manner. To maximize our statistical power to identify such differences we chose to test this hypothesis in a disease setting with large number of relatively homogenous cell lines and fewer unknown covariates. Using CanDI, we stratified all KRAS-mutant NSCLC, pancreatic adenocarcinoma (PDAC), and colorectal cancer (CRC) by sex and then tested for conditional gene essentiality. This analysis identified a number of genes that are differentially essential in male or female KRAS-mutant NSCLC, PDAC and CRC models (Fig. 3a-f and Supplementary Table 5). The genes that we identify are not common across all three disease types suggesting as one might expect that the biology of the tumor in part also determines gene essentiality. To test whether any association between differentially essential genes could be identified from expression data (e.g essential genes encoded on the Y chromosome) we first used CanDI to identify genes that are differentially expressed between male and female cell lines within each disease ^31^. We then plotted the set of differentially essential genes against the differentially expressed genes in KRAS-mutant NSCLC, PDAC and CRC models (Fig. 3a,c,e and Supplementary Table 6) and found little overlap between these gene lists. A number of genes that are more essential in male cells, such as *AHCYL1, ENO1, GPI* and *PKM*, regulate cellular metabolism. This finding is consistent with previous literature on sex and metabolism^32^. Our analysis demonstrates that stratifying groups of heterogeneous cancer models by three variables, in this case tumor type, KRAS mutation status and sex, reveals differentially essential genes. CanDi enables biologically principled stratification of data in the CCLE and DepMap by any feature associated with a group of cell models. This stratification allows us to identify genes associated with sex, which is not possible with other covariates included.

**Figure 3.**
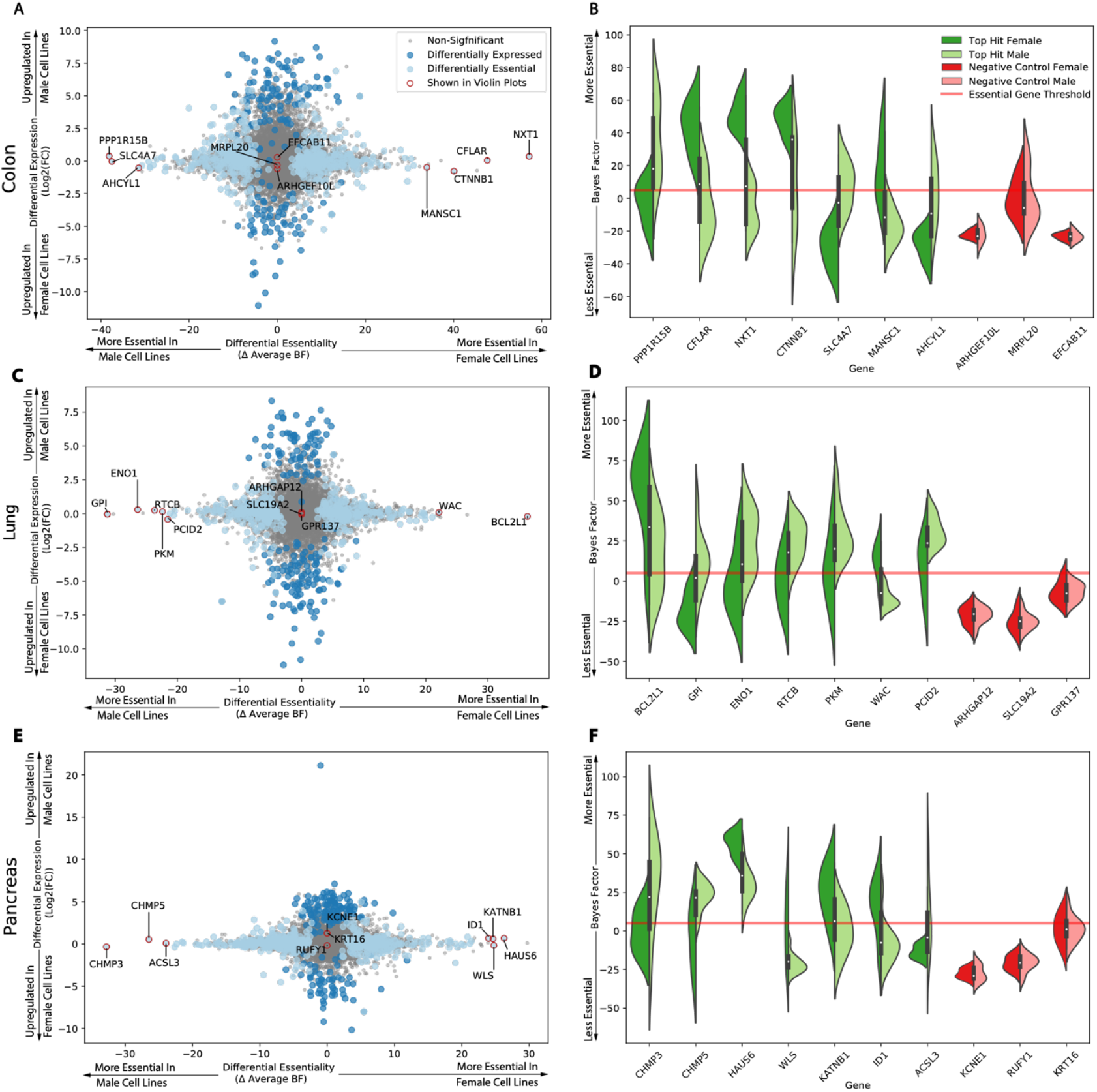
**(A)** Differential gene expression and differential gene essentiality in male and female CRC cell lines. N=7 male cell lines and N=3 female cell lines. **(B)** The distribution of Bayes factor gene essentiality scores in male and female CRC cell lines. The top seven and bottom three differentially essential genes are shown in violin plots split by the sex of the cell lines. **(C)** Differential gene expression and differential gene essentiality in male and female NSCLC cell lines. N=9 male cell lines and N=5 female cell lines. **(D)** The distribution of Bayes factor gene essentiality scores in male and female NSCLC cell lines. The top seven and bottom three differentially essential genes are shown in violin plots split by the sex of the cell lines. **(E)** Differential gene expression and differential gene essentiality in male and female PDAC cancer cell lines. N=13 male cell lines and N=5 female cell lines. **(F)** The distribution of Bayes factor gene essentiality scores in male and female PDAC cell lines. The top seven and bottom three differentially essential genes are shown in violin plots split by the sex of the cell lines.

### CanDI enables rapid integration of external datasets to reveal new immunotherapy targets

An emerging challenge in the cancer biology is how to robustly integrate larger “resource” datasets like CCLE with the vast amount of published data from individual laboratories. For example, a big challenge in antibody discovery is identifying specific surface markers on cancer cells. To approach these big questions we utilized CanDIs ability to rapidly take new datasets, such as raw RNA-seq counts data in a disparate study of interest, then normalize and integrate this data into the CCLE, DepMap and protein localization databases previously described. Specifically, we rapidly integrated an RNA-seq expression dataset that measured the set of transcribed genes in primary lung bronchial epithelial cells from 4 donors ^33^. Classes within CanDI enable rapid application of DESeq2 to assess the differential expression between outside datasets and the CCLE. We used this feature to identify genes that are differentially expressed between primary lung bronchial epithelial cells and KRAS-mutant NSCLC, EGFR-mutant NSCLC or all NSCLC models in CCLE. We then used CanDI to identify genes that are upregulated in cancer cells over normal lung bronchial epithelial cells with protein products that are localized to the cell membrane. This analysis of KRAS-mutant, EGFR-mutant and pan-NSCLC generated highly similar lists of differentially expressed surface proteins (Fig. 4a-f and Supplementary Table 7). Notably, overexpression of several of these genes, such as *CD151* and *CD44*, has been observed in lung cancer and is associated with poor prognosis ^34–36^. These proteins represent potential new immunotherapy targets in KRAS-driven NSCLC.

**Figure 4.**
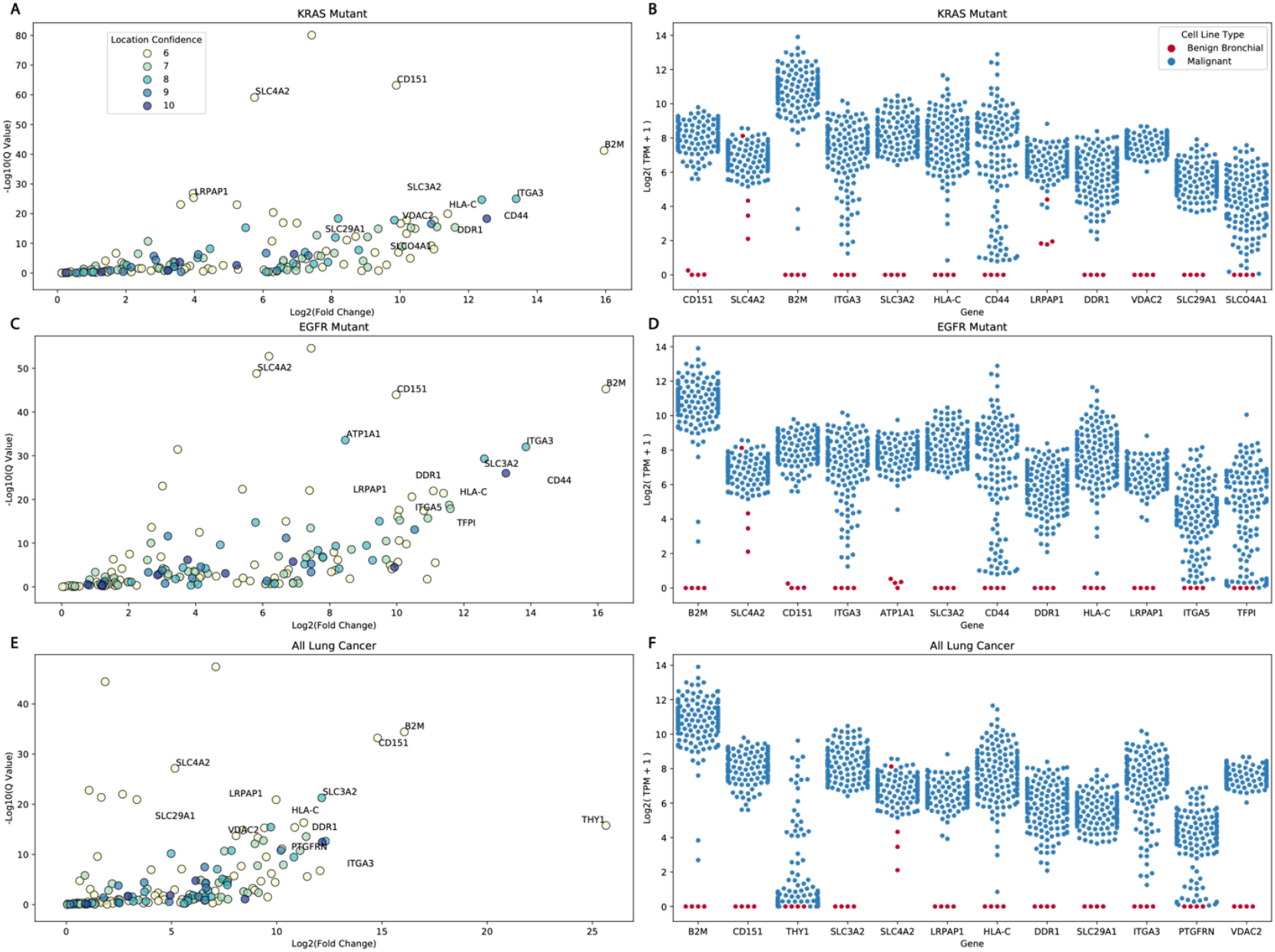
**(A)** A graph showing genes that are upregulated in KRAS-mutant NSCLC cell lines relative to primary human bronchial epithelial cells. A cell membrane protein localization score is shown for each gene. Higher protein localization scores indicate higher confidence annotations. **(B)** A scatter plot showing gene expression for genes that encode cell surface proteins in KRAS-mutant NSCLC cell lines and primary human bronchial epithelial cells. N=46 for KRAS-mutant NSCLC cell lines and N=4 for primary human bronchial epithelial cells. **(C)** A graph showing genes that are upregulated in EGFR-mutant NSCLC cell lines relative to primary human bronchial epithelial cells. A cell membrane protein localization score is shown for each gene. Higher protein localization scores indicate higher confidence annotations. **(D)** A scatter plot showing gene expression for genes that encode cell surface proteins in EGFR-mutant NSCLC cell lines and primary human bronchial epithelial cells. N=21 for EGFR-mutant NSCLC cell lines and N=4 for primary human bronchial epithelial cells. **(E)** A graph showing genes that are upregulated in NSCLC cell lines relative to primary human bronchial epithelial cells. A cell membrane protein localization score is shown for each gene. Higher protein localization scores indicate higher confidence annotations. **(F)** A scatter plot showing gene expression for genes that encode cell surface proteins in NSCLC cell lines and primary human bronchial epithelial cells. N=141 for NSCLC cell lines and N=4 for primary human bronchial epithelial cells.

## Discussion

Data integration is a critical requirement in biology research in the era of genomics and functional genomics. Large scale efforts such as the CCLE have revealed genomic features of more than 1000 cell line models. This data has not to our knowledge previously been integrated with functional genomics data in a manner that individual users can enter batched queries that are stratified by disease subtype or mutation status. This is not just a small improvement in functionality, but rather it is an enabling format that makes possible the types of conditional genomics analyses that drive discovery. Moreover, it fills a fundamental gap in the cancer research community that integrates large scale projects with investigator initiated studies Our data framework enables biologists without specialized expertise in bioinformatics to use the full spectrum of data in the CCLE and DepMap in a higher throughput and precise manner. Using CanDI, we identified genes that are selectively essential in male versus female KRAS-mutant NSCLC, PDAC and CRC models. To our knowledge, such analysis has never been performed to begin to query the biologic basis of sex disparity in cancer or cancer therapy. We illustrate another feature of our framework by analyzing a list of hit genes nominated by a bespoke CRISPR drug screen for gene essentiality in BRCA1/2-wild type and BRCA1/2-mutated breast and ovarian cancer. In a third application, we analyzed the principle of synthetic lethality for 17427 genes in 19 KRAS-mutant and 11 EGFR-mutant NSCLC models. We then used CanDI to globally identify genes that are conditionally essential in the context of common cancer driver mutations. Finally, we nominated 12 potential new immunotherapy targets in KRAS-mutant, EGFR-mutant and pan -NSCLC models by using CanDI to identify genes that are differentially expressed in normal bronchial epithelial cells versus NSCLC models that are localized at the plasma membrane. Our data reveal a wealth of new hypotheses that can be rapidly generated from publicly available cancer data. By sharing data flows and use cases with a CanDI community we illustrate the ways in which individual research groups can interact with massive cancer genomics projects without reinventing tools or relying upon DepMap tool releases. We anticipate that CanDI will be widely used in cell biology, immunology and cancer research.

## Methods

### CanDI

The CanDI data integrator is available at https://github.com/Yogiski/CanDI.

### CanDI Module Structure

The CanDI data integrator is a python library built on top of the Pandas that is specialized in integrating the publicly available data from The Cancer Dependency Map (DepMap Release: 2019 Quarter 3)^12^, The Cancer Cell Line Encyclopedia (CCLE Release: 2019 Quarter 3) ^1^, The Pooled In-Vitro CRISPR Knockout Essentiality Screens Database (PICKLES Library: Avana 2018 Quarter 4) ^20^, The Comprehensive Resource of Mammalian Protein Complexes (CORUM)^8^ and protein localization data from The Cell Atlas^4^, The Map of the Cell^11^, and The In Silico Surfaceome^7,21^. Data from DepMap and CCLE used in the following analyses are from the 2019Q3 release. Data from PICKLES is from the 2018 Quarter 4 release of DepMap using the Avana library.

Access to all datasets is controlled via a python class called Data. Upon import the data class reads the config file established during installation and defines unique paths to each dataset and automatically loads the cell line index table and the gene index table. Installation of CanDI, configuration, and data retrieval is handled by a manager class that is accessed indirectly through installation scripts and the Data class. Interactions with this data are controlled through a parent Entity class and several handlers. The biologically relevant abstraction classes (Gene, CellLine Cancer, Organelle, GeneCluster, CellLineCluster) inherit their methods from Entity. Entity methods are wrappers for hidden data handler classes who perform specific transformations, such as data indexing and high throughput filtering.

### Differential Expression

In all cases where it is mentioned differential expression was evaluated using the DESeq2 R package (Release 3.10) ^31^. Significance was considered to be an adjusted p-value of less than 0.01.

### Differential Essentiality

Essentiality scores are taken from the PICKLES database (Avana 2018Q4). To reduce the number of hypotheses posed during this analysis the mutual information of gene essentiality was calculated using the mutual information metric from the python package SciKitLearn (Version 0.22.0). Genes with mutual information scores greater than one standard devation above the median were removed from consideration. Differential essentiality was evaluated by performing a Mann-Whitney u-test between two groups on every gene that passed the mutual information filter. Significance was considered to be a p-value of less than 0.01. Magnitude of differential essentiality of a given gene was shown as the difference in mean Bayes factors between two groups of cell lines.

### Protein Localization Confidence

Protein localization data was assembled from The Cell Atlas^4^, The Map of the Cell^11^, and The In Silico Surfaceome^7,21^. Confidence annotations were taken from the supplemental data of each paper and put on a number scale from 0 to 4 and summed for a total confidence score for each localization annotation for every gene where across all three papers. The analysis shown in Figure 4 represents a gene list that was further manually curated to remove the genes that are localized to the intracellular space at the cell membrane revealing cell surface protein targets that are highly expressed in NSCLC cancer models over normal lung bronchial epithelial cells ^4,7,11,21^.

### DepMap Creative Commons License

When an individual user runs CanDI they are downloading DepMap data and thus are agreeing to a <u>CC Attribution 4.0 license</u> (https://creativecommons.org/licenses/by/4.0/).

### Synthetic Lethality of Fanconi Anemia Genes in Ovarian and Breast Cancer Models

We made a list of the top 50 gene hits that confer sensitivity to PARP inhibition in HeLa cells^23^. Using CanDI the essentiality scores of these top hits were visualized across all ovarian cancer cell models in PICKLES (Avana 2018Q4). FANCA and FANCE showed selective essentiality in the BRCA1/2 mutant ovarian cancer cell lines. Following this observation CanDI was used to gather the gene essentiality for all FANC genes in the fanconi anemia pathway. CanDI was then used to visualize these data across all ovarian and breast cancer cell lines, sorting by BRCA1/2 mutation status.

### Synthetic Lethality in KRAS and EGFR mutant Cell Lines

CanDI was leveraged to bin NSCLC cell lines present in both CCLE (Release: 2019Q3) and PICKLES (Avana 2018Q4) into 8 groups. KRAS mutant and KRAS wild type cell lines with and without EGFR mutants removed as well as EGFR mutant and EGFR wild type cell lines with and without KRAS mutants removed. The mean essentiality score for every gene in the genome was calculated for every group of cell lines. Synthetic lethality score per gene is defined as the change in mean essentiality from the mutant groups to the wild type groups.

### Pan Cancer Synthetic Lethality Analysis

A set of 299 core oncogenes and tumor suppressor driver mutations was chosen for analysis^37^. To test the effect of these gene’s mutations on gene essentiality CanDI was leveraged to split into two groups: a nonsense mutation group containing genes annotated as tumor suppressors (N=153) and a missense mutation group containing genes annotated as oncogenes with specific driver protein changes (N=53). CanDI was then used to collect a core set of genes with highly variable essentiality. To do this the Bayes factors from the PICKLES database (Avana 2018Q4) were converted to binary numeric variables. Bayes factors over 5 were assigned a 1=essential and Bayes factors under 5 were assigned a 0=non-essential. Genes were then sorted buy their variance across cell lines and genes between the 85^th^ and 95^th^ percentile were used for this analysis (N=2340). To determine a short list of genes with which to follow up on Chi^2^ tests were applied to the 95940 gene pairs in the missense group and the 603720 gene pairs in the tumor suppressor group. Three new groups were formed for further analysis: the first consisted of the significant gene/mutation pairs from the oncogenic group, the second consisted of the significant gene/mutation pairs from the tumor suppressor group, and the third was a combination of the significant pairs from both groups with no discrimination on the type of mutations considered.

These groups were further analyzed for differential essentiality via the Mann Whitney method described above and the Cohens D effect size were calculated to measure the extent of the phenotype.

### Differential Expression and Essentiality of Male and Female KRAS driven cancers

We used CanDI to gather all cell lines that are present in both PICKLES (Avana 2018Q4) and CCLE (Release 2019Q3). CanDI was then leveraged to put these cell lines into the following tissue groups: KRAS mutant Colon/Colorectal, PDAC, and NSCLC. Each tissue group was then split into male and female sub-groups. Differential expression was analyzed by applying the methods described above to raw RNA-seq counts data from CCLE (Release: 2019Q3). Genes with adjusted p-values less than 0.01 were considered significantly differentially expressed.

Differential essentiality was analyzed using the methods described above on the previously described sex-subgroups for each tissue type. Genes with p-values less than 0.01 were considered significantly differentially essential between male and female cell models. For each tissue type the distributions of the top 7 significantly differentially essential genes were highlighted in comparison with the bottom 3 as a negative control.

### Differential expression of benign and malignant cancer cell lines

We downloaded human bronchial epithelial (HBE) RNA-seq data from Gillen et al via the European Nucleotide Archive to use as a benign lung tissue model^33^. This 4 data set contains gene expression data for primary HBE cells cultured from three different donors and also NHBE cells (Lonza CC-2541, a mixture of HBE and human tracheal epithelial cells). We then used CanDI to put NSCLC models into three different groups: KRAS mutant, EGFR mutant, and all cell lines. For our benign model raw counts were quantified via kallisto^38^. Raw counts for our malignant cell lines were queried via CanDI. DESeq2 was then applied to evaluate the differential expression between our normal lung tissue model and our three malignant lung tissue groups. The results from DESeq2 were then filtered by significance (adjusted p-value < 0.01). To filter based on potential immunotherapy targets we removed all genes not annotated as being localized to the plasma membrane, and genes with localization confidence scores lower than six. Genes that were obviously mis-annotated as surface proteins were also manually removed.

## Supporting information

Zip with all supplementary data

## Supplementary Figure/Table Legends

**Supplementary Figure 1.**
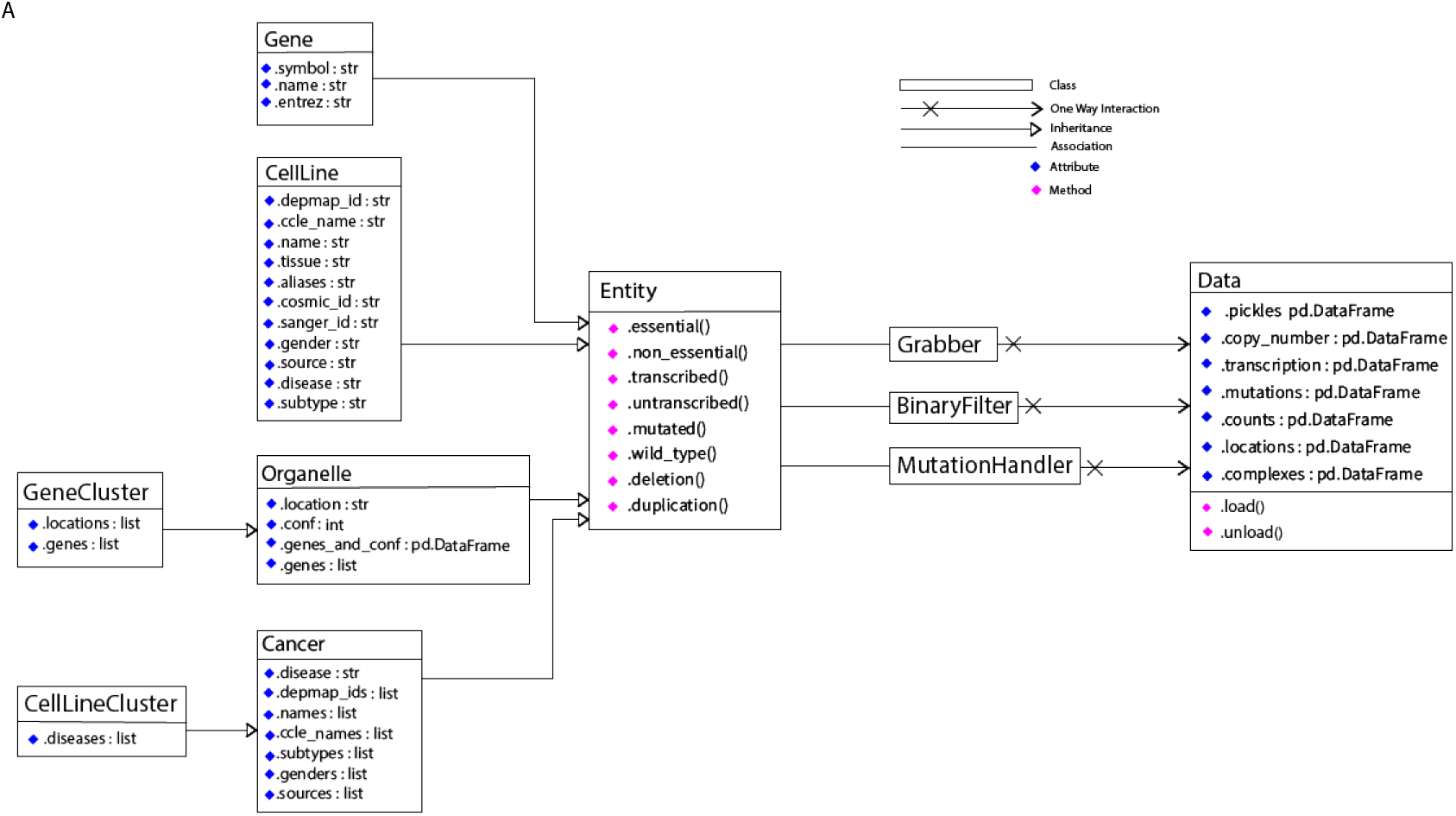
An Object-oriented schema diagram showing core structure of CanDI software.

**Supplementary Table 1**. A table containing raw PICKLES Bayes factors displayed in the heat map of Fig. 1e.

**Supplementary Table 2**. A table containing mean PICKLES Bayes factors for each series displayed in Fig. 2a,b.

**Supplementary Table 3**. A table containing the data for all chi^2^ tests performed to generate Fig. 2c,d.

**Supplementary Table 4**. A table containing the data for scatter plots shown in Fig. 2e,f,g. **Supplementary Table 5**. A table containing the data from the differential essentiality analysis for all three tissues in Fig. 3a-f.

**Supplementary Table 6**. A table containing the data from the differential expression analysis for all three tissues in Fig. 3a,c,e.

**Supplementary Table 7**. A table containing the differential expression analysis data merged with the location data for all three tissues shown in Fig. 4.

## Acknowledgements

We thank everyone in the Gilbert lab for helpful comments and discussion. LAG is supported by K99/R00 CA204602 and DP2 CA239597 as well as the Goldberg-Benioff Endowed Professorship in Prostate Cancer Translational Biology.

